# dynUGENE: an R package for uncertainty-aware gene regulatory network inference, simulation, and visualization

**DOI:** 10.1101/2021.01.07.425782

**Authors:** Tianyu Lu, Anjali Silva

## Abstract

Methods for gene regulatory network inference focus on network architecture identification but neglect model selection and simulation. We implement an extension to the dynGENIE3 algorithm that accounts for model uncertainty as an R package, providing users with an easy to use interface for model selection and gene expression profile simulation. Source code is available at https://github.com/tianyu-lu/dynUGENE with a detailed user guide. A webserver with interactive controls is available at https://tianyulu.shinyapps.io/dynUGENE/.

## Introduction

Complex phenomena such as cell development and apoptosis emerge from coordinated dynamics of gene regulatory networks (GRN). Inferring network structure from data can be used for hypothesis generation, revealing mechanisms in cell development and disease (Huang et al., 2009), and modelling network evolution (Crombach and Hogeweg, 2008). Accurate dynamical models allow us to predict the effects of network perturbations on biological function, for example to push cells out of a disease state (Karlebach and Shamir, 2010), or to design synthetic GRNs given the desired dynamics of a network (Hiscock, 2019). The ideal model should be flexible enough to capture highly nonlinear interactions while not sacrificing model interpretability and computation time.

We present dynUGENE (dynamical Uncertainty-aware GEne NEwork inference), an R package that extends the functionality of dynGENIE3, a state-of-the-art method for GRN inference (Geurts et al., 2018). We build on dynGENIE3 because it satisfies all three of our model desiderata. Existing extensions include TIMEOR and BENIN which both incorporate heterogeneous data to improve network inference accuracy (Wonkap and Butler, 2020; Conard et al., 2020). Here, we take a different approach and instead account for uncertainty in dynGENIE3, allowing for stochastic gene expression simulations and parsimonious model selection. Our extension is available as an easy to use R package and also as an interactive web server.

## Package Design

### dynGENIE3 Background

dynGENIE3 poses GRN inference as a feature selection problem. It first trains random forests to predict the change in concentration of each species given the current concentrations of all species. Each interaction from species *x_i_* to species *x_j_* is associated with an importance score, calculated by the reduction in variance from using *x_i_* to predict the change in *X_j_*. The importance score for an interaction, when normalized, is interpreted as the probability of that interaction to exist. For a detailed treatment, see the vignette and (Geurts et al., 2018).

### Model Selection

The inferred network can be visualized as a *p* × *p* matrix where the entry [*x_i_, x_j_*] is the importance score of *x_i_* for inferring *x_j_* (Fig. 1). However, real GRNs are often not fully connected and the presence of an interaction is binary (Mangan et al., 2016). To address this, dynUGENE includes a function for model selection based on visualizing the Pareto front (Mangan et al., 2016). However, we note that the model at the sharp drop in the Pareto front is not always the best model (Supplementary Fig. S1). We include an additional function on the web server where users can choose which interactions to mask. The masked networks can then be simulated, allowing for application-specific tuning of model complexity.

**Fig. 1:**
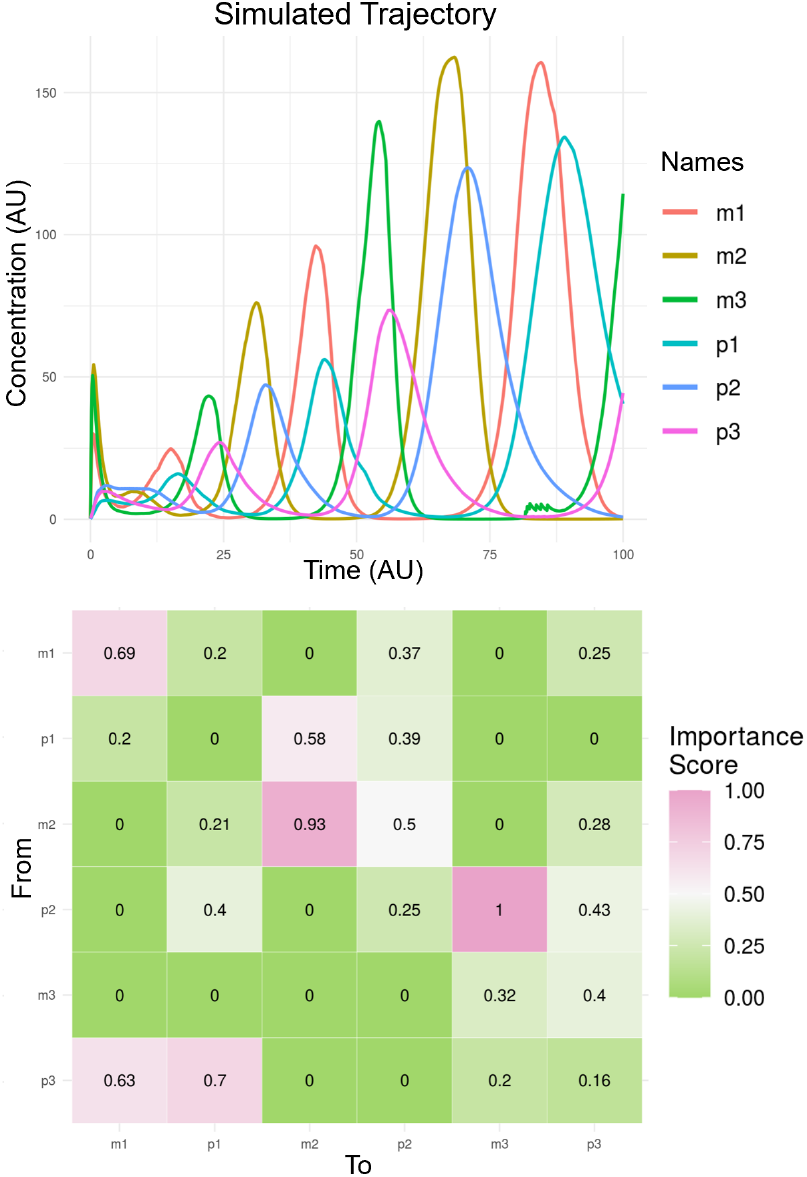
Bottom: inferred importance scores on the repressilator dataset for the 16^th^ network in the step-wise column masks plot (Supplementary Fig. S2). Top: Simulated trajectory using the inferred network.

### Model Simulation

The inferred networks and masked networks can be used to simulate gene expression profiles by numerically solving the system of ordinary differential equations learned by the random forests. In addition to deterministic simulations, we provide an option that accounts for the uncertainty in the random forests predictions for stochastic simulations. For stochastic simulations, instead of only taking the mean of a random forest’s predictions, we sample from the Gaussian 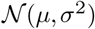 where *μ* is the mean and *σ*^2^ is the variance of the random forest’s predictions.

### Provided Datasets

The dynUGENE package provides four example time-series datasets: repressilator, stochastic re-pressilator, Hodgkin-Huxley, and stochastic Hodgkin-Huxley (Elowitz and Leibler, 2000; Hodgkin and Huxley, 1952). These datasets were generated from systems of ordinary or stochastic differential equations. Details are provided in the vignette. The package also includes one steady state dataset, SynTReN3OO, taken from GRNdata (Bellot et al., 2020). Users can provide their own data as input following the for-mat specified in `?inferNetwork`.

## Discussion

A requirement for dynGENIE3 and dynUGENE is that all species must be tracked through time. This requirement is difficult to satisfy in practice as there are often unknown species in a biological process of interest. Methods that can identify or approximate latent structure in partially-observed systems are more appropriate here (Hiscock, 2019). An omics treatment such as RNA-seq can cover breadth but current sequencing techniques require cells to be destroyed, thus making time series data collection difficult. Non-destructive sequencing techniques could address this issue.

The implementation of an inferred network as a gene circuit will require more thought. Even for networks with sparse interactions, the likelihood of finding a set of genes and proteins that satisfy the interaction strengths and activation or inhibitory effects is unknown. In fact, whether a species is an activator or inhibitor is not explicitly given in the interaction matrix. We can address this by posing dynUGENE as a constrained optimization problem where it is limited to using only a given set of parts (genes, promoters, ribosome binding sites, proteins, etc.) thus relating the importance scores with biological interaction strengths. We leave this for future work.

## Data and code availability

Source code is available at https://github.com/tianyu-lu/dynUGENE with a detailed user guide. A webserver with interactive controls is available at https://tianyulu.shinyapps.io/dynUGENE/.

## ACKNOWLEDGEMENTS

The authors thank the authors of dynGENIE3 for their work and Alan Moses for guidance.

## FUNDING

This work was supported by a Postdoctoral Fellowship from Canadian Institutes of Health Research.

## Notes

https://github.com/tianyu-lu/dynUGENE

https://tianyulu.shinyapps.io/dynUGENE/

### Competing Interest Statement

The authors have declared no competing interest.

