## Supplemental Figures for "dynUGENE: an R package for uncertainty-aware gene regulatory network inference, simulation, and visualization"

### Supplementary Information

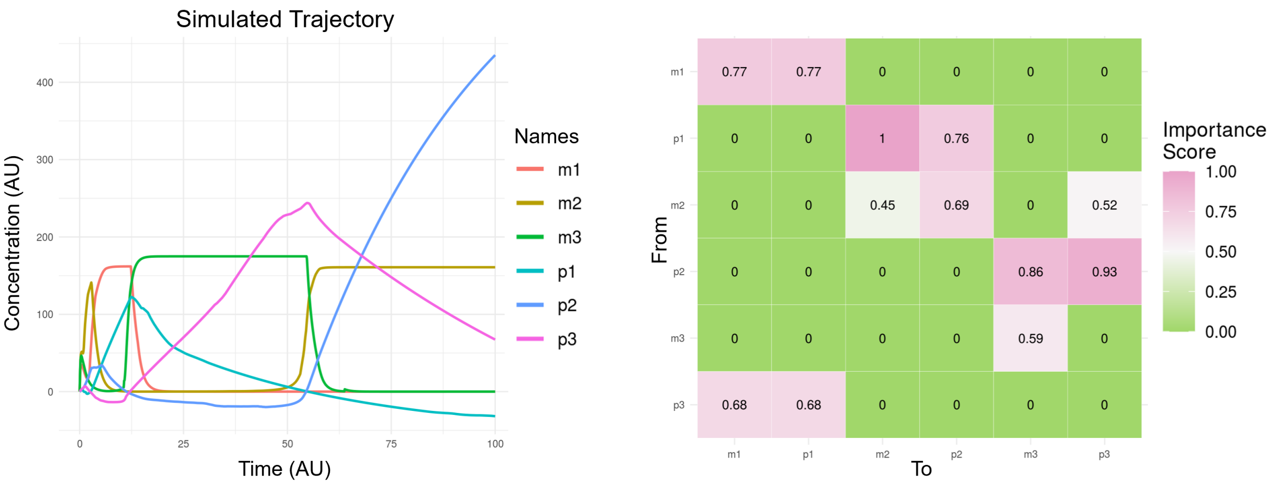


**Supplementary Fig. S1:** Left: Simulated trajectory using the inferred network. Right: inferred importance scores on the repressilator dataset for the $7^{th}$ network in the step-wise column masks plot (Supplementary Fig. S2).

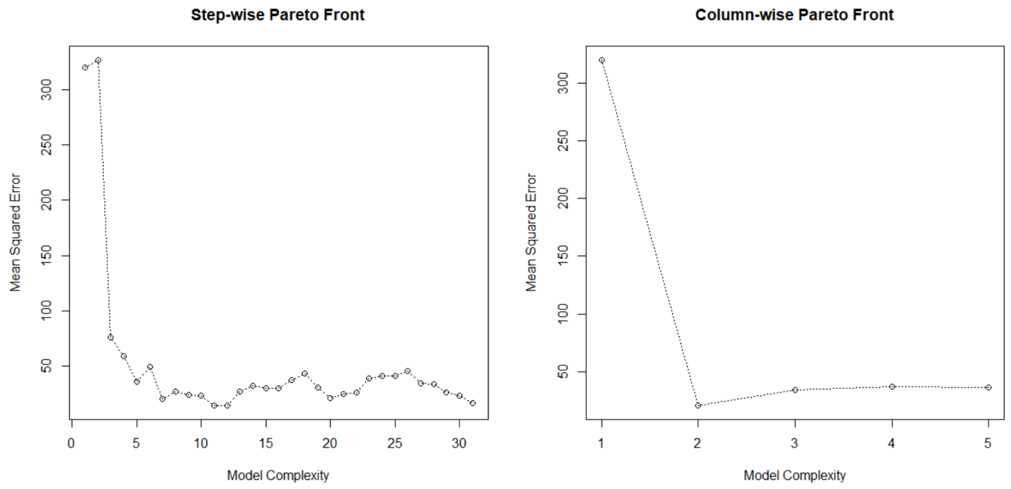


**Supplementary Fig. S2:** Prediction error vs. model complexity for the repressilator dataset. The simplest network for step-wise masks only keeps the interactions within the highest importance scores per column of the interaction matrix. Model complexity increases by including the interaction with the next highest importance score. For column-wise masks, model complexity increases by including the interactions with the next highest importance scores, one per column.
